# Increased CPT1a expression is a critical cardioprotective response to pathological stress that suppresses gene programs for remodeling and enables rescue by gene transfer

**DOI:** 10.1101/2024.09.30.615965

**Authors:** Andrew N. Carley, Santosh K. Maurya, Chandan K. Maurya, Yang Wang, Amy Webb, Azariyas A. Challa, Zhentao Zhang, Hua Zhu, Ahlke Heydemann, Kenneth C. Bedi, Christos P. Kyriakopoulos, Craig H. Selzman, Stavros G. Drakos, Kenneth B Margulies, E. Douglas Lewandowski

**Author notes:** **Corresponding Author:** Douglas Lewandowski, Ph.D. The Ohio State University Medical Center Biomedical Research Tower, BRT Rm 312 460 W 12^th^ Avenue Columbus, OH 43210.

## Abstract

**Background:** Carnitine palmitoyl transferase 1 (CPT1) is a rate-limiting enzyme for long chain fatty acid oxidation (FAO) in cardiac mitochondria. In adult hearts, CPT1b predominates, while CPT1a is co-expressed at lower levels. Pathological stress on the heart induces greater CPT1a expression, and this coincides with a reduction in FAO, yet the role of CPT1a in pathological cardiac remodeling is unknown.

**Methods:** CPT1 isoform expression was assayed in myocardium of human heart failure (HF) patients with nonischemic cardiomyopathy (NICM) and a preclinical mouse model of heart failure. To explore the role of CPT1a upregulation in response to pathological stress, mice were subjected to afterload stress via transverse aortic constriction (TAC) or sham surgery (sham) with cardiac-specific CPT1a knockdown or cardiac-specific, AAV9-mediated CPT1a overexpression (AAV9.cTNTN.Cpt1a), versus empty virus or PBS infusions as controls. MiR370, known to suppress hepatic CPT1a, was assayed and overexpressed to determine if miR370 regulates cardiac CPT1a expression.

**Results:** CPT1a protein was elevated and miR370 reduced in myocardium of male and female NICM patients (204% vs. non-failing unused donor hearts), as well as in failing mouse hearts. AAV mediated miR370 overexpression in mouse hearts suppressed CPT1a expression and attenuated the response of CPT1a to TAC. Preventing CPT1a upregulation in response to TAC in cardiac specific CPT1a knockout mice (csCPT1a ko) exacerbated adverse remodeling, causing severe dysfunction and increased mortality. In contrast, CPT1a overexpression (2.8 fold), attenuated impaired ejection fraction (EF, by 54%) and fractional shortening (FS, 65%) vs. PBS-infused TAC hearts (p<0.05). Delivery of AAV9.cTnT.Cpt1a 4 wks after TAC surgery, led to significant rescue of EF and FS vs. animals receiving empty virus and mitigated the exacerbated dysfunction of csCPT1a ko hearts at 4 wks TAC. RNA-seq and reverse transcription-quantitative PCR revealed a novel function of CPT1a in suppressing hypertrophic, profibrotic and cell death gene programs in both sham and TAC hearts, irrespective of changes in FAO.

**Conclusions:** The effects of CPT1a in the heart extend beyond FAO and include a non-canonical regulation of cardiac gene programs. In addition to an animal model of HF, CPT1a upregulation occurs in NICM, and is a critical cardioprotective adaptation to pathological stress.

## Introduction

In response to pathological stress, the adult heart upregulates several fetal isoform genes ^1,2^, including a shift in the relative expression of two co-expressed isoforms of the rate limiting enzyme for long chain fatty acid oxidation (FAO) in cardiomyocytes, carnitine palmitoyl transferase 1 (CPT1) ^3–5^. In adult hearts, the CPT1b isoform is in greater abundance than CPT1a, which is more highly expressed in the fetal heart ^3–5^, each having distinct enzyme kinetics and different modes of allosteric regulation ^4^. An early component of the response to pathological stress on the heart in animal models is increased CPT1a expression prior to cardiac dysfunction ^6^ and persisting into overt heart failure (HF) ^3–5,7^. The role of CPT1a expression in the cardiac response to pathological stress is to date unexplored.

This increase in CPT1a coincides with the well-reported reduction in FAO and increased reliance, albeit inefficient, on glucose for ATP production ^8,9^, a pattern of substrate utilization consistent with human HF with reduced ejection fraction (HFrEF) ^10,11^. Elevating CPT1a content by cardiac-specific gene delivery in otherwise normal hearts results in a paradoxical reduction in FAO that recapitulates the phenotype of pathological hearts ^12^. These effects of CPT1a on cardiac FAO occur in the absence of changes in CPT1b. Although CPT1a has been shown to be less sensitive to inhibition by malonyl CoA than CPT1b in isolated mitochondria from liver and muscle cells ^13^, and could be expected to facilitate higher rates of FAO, this is not the case in the heart ^12,14–16^. Furthermore, augmenting CPT1a in healthy rodent hearts induces cardiac natriuretic peptide (NP) expression, as also occurs in response to afterload stress when CPT1a becomes upregulated ^12^. Thus, CPT1a, and potentially upstream factors controlling CPT1a expression, appears to provide a fundamental link between metabolic remodeling and cardiac pathology. Whether this shift toward elevated CPT1a is adaptive or maladaptive is entirely unknown.

Therefore, we examined CPT1a content and that of a microRNA known to control hepatic CPT1a expression, miR370 ^17^, in the left ventricular myocardium of patients with non-ischemic cardiomyopathy (NICM) and the metabolic and pathophysiological consequences of altered CPT1a expression in mouse models of afterload stress. Cardiac specific CPT1a genetic knockdown, adeno-associated virus serotype 9 (AAV9)-mediated, cardiac specific CPT1a overexpression, and also (AAV9) mediated cardiac specific, miR370 overexpression were all explored.

Results show for the first time increased CPT1a protein in hearts of HFrEF patients, replicated in two different NICM patient cohorts at two different institutions and that induction of CPT1a is an adaptive rather than maladaptive stress response that attenuates adverse cardiac remodeling. Further, we have elucidated a directional regulation of CPT1a expression by miR370 in a preclinical animal model, which correlates to changes in CPT1a and miR370 in human HF, and a direct effect of CPT1a expression in mediating cardiac NP production. Beyond the role of CPT1a in fat metabolism, we report the first evidence that CPT1a functions in adult myocardium to inhibit gene programs associated with adverse cardiac remodeling, including profibrotic, hypertrophic, and cell death responses. The findings reveal CPT1a is requisite in the adaptive response to pathological stress, independently of FAO to attenuate contractile dysfunction in the failing heart, and that CPT1a overexpression confers cardioprotection.

## Methods

Data, analytic methods, and study materials will be made available to researchers for purposes of reproducing results or replicating procedures upon reasonable request. Full expanded methodological details are provided in the Supplemental Material. Wherever possible researchers were blinded to sample identity for assays and analysis.

### Animals

Cardiac specific knockdown of CPT1a (csCPT1a ko) (n=57) was generated by crossing CPT1a floxed (f/f) mice (see online supplement) with hemizygous *α*MHC-Cre mice (stock# 011038, The Jackson Laboratory). Control mice were f/f littermates (n=56).

CPT1a was overexpressed in 10-12 week old male C57Bl/6 mice (n=71) (The Jackson Laboratory) with adeno-associated virus (AAV) 9 delivery under control of the cardiac specific promoter cTNT via jugular vein injection (AAV9.cTNT.CPT1a) (Virovek, www.virovek.com). To suppress CPT1a, miR370 (n=11) was inserted into the same AAV9 construct and injected i.v. Table S1 and Figure S1 show vector design and sequences. Control mice received either PBS (n=46) or AAV9 containing an empty vector sequence (AAV9.Emp) (n=24).

As a similar increase in CPT1a occurred in failing hearts of male and female NICM patients, only male mice were studied. Mice were randomly distributed to experimental protocols, housed in the same location, without exclusions. All experimental procedures involving vertebrate animals were approved by the Institutional Animal Care and Use Committees (IACUC) at the University of Illinois at Chicago and the Ohio State University.

### Pathological cardiac hypertrophy

Pathological hypertrophy was induced by transverse aortic constriction (TAC), as previously described ^18^ with modification for different clip sizes. CsCPT1a ko mice and f/f littermate controls underwent TAC using a 0.018 inch micro-clip or sham surgery (sham) at 10-14 weeks of age. A subset of csCPT1a ko mice underwent TAC surgery (0.018 clip) and 1 wk later received AAV9.cTnT.Cpt1a or AAV9.Emp.

Mice injected with AAV9.cTnT.miR370 or AAV9.cTnT.Cpt1a (i.v.) underwent TAC 10 d after injection with a 0.016 inch micro-clip. A subset of C57Bl/6 mice underwent TAC (0.016 clip) or sham surgery followed by AAV9.cTnT.Cpt1a or AAV9.Emp 4 wks post-TAC.

Transthoracic ultrasound imaging was performed to track cardiac hypertrophy and functional remodeling (Vevo 2100, VisualSonics) ^18^.

### Isolated heart perfusion

Eight weeks post-TAC or sham, mice were randomly selected and hearts isolated for retrograde perfusion with modified Krebs-Henseleit buffer containing 0.4 mmol/L unlabeled palmitate/albumin complex (3:1 molar ratio), 10 mmol/L glucose, and 1 mmol/L lactate ^19^. Prior to cardiectomy, mice received heparin (50 U/10 g) and anesthesia (80 mg/kg Ketamine and 12 mg/kg xylazine, i.p.). Hearts were situated in a 10 mm broadband probe within a vertical wide-bore (89mm) 14.1 T NMR magnet. Sample chamber temperature was maintained at 37 °C. Initially, a two-minute ^31^P NMR spectrum was acquired, followed by a natural abundance ^13^C spectrum for subtraction of endogenous background signal. Perfusate was switched to similar media containing [2,4,5,6,8,10,12,14,16-^13^C_8_] palmitate for 30 min ^19^. Hearts were frozen in liquid N_2_ cooled tongs.

#### In vitro NMR spectroscopy for substrate selection

The fractional contribution (Fc) of ^13^C palmitate into the tricarboxylic acid (TCA) cycle as acetyl CoA was determined by glutamate isotopomer analysis from *in vitro* ^13^C NMR spectra of left ventricular tissue extracts ^20^.

### Human heart tissue collection

Protocols for tissue sampling at the time of heart transplantation were approved by the University of Pennsylvania Institutional Review Board and for transmural LV apical core sampling at CF-LVAD implant by the University of Utah Institutional Review Board. Written informed consent for use of heart tissues was obtained prospectively from transplant recipient patients and from next-of-kin in the case of organ donors. Use of hearts from brain-dead organ donors for sampling at the University of Pennsylvania (n=8) was approved by the Gift-of-Life donor Program in Philadelphia, PA. Use of non-failing donor hearts not allocated for human transplantation because of noncardiac reasons were approved for use at the University of Utah (n=9) by DonorConnect of Salt Lake City, UT. We selected individuals with advanced HF due to non-ischemic cardiomyopathy (NICM). All samples were from subjects without history of diabetes. Procurement of human myocardial tissue (n=8) at the University of Pennsylvania was performed as previously described ^21,22^. Transmural apical biopsies at University of Utah (n=9) were immediately frozen and stored at -80°C for metabolic enzyme expression analysis. See Tables S2 And S3 for details and patient and donor characteristics.

### Metabolic enzyme expression and content

Protein expression was determined by western blot by loading 20 ug of protein per lane and band intensities quantified by LI-COR Odyssey Fc or BioRad ChemiDoc and normalized to the expression of calsequestrin (CASQ) or glyceraldehyde-3-phosphate dehydrogenase (GAPDH) as loading controls. mRNA levels were determined by quantitative reverse transcription polymerase chain reaction in frozen heart tissue and normalized to S29 ^18^. Antibodies, antibody dilutions (Table S4) and primers (Table S5) are listed in the Supplement.

### MiR370 Assay

MiR370 expression was measured using the Taqman MicroRNA assays, 462611_mat for mice and assay ID 002275 Cat# 4427975 for human samples (Applied Biosystems) and normalized to U6 expression (Assay ID 001973, Cat# 4427975).

### Immunohistochemistry on heart sections

Immunohistochemistry was performed on hearts 10d post AAV.Cpt1a or AAV9.Emp injection to access phospho-histone H3 levels.

### RNA-Seq

RNA-Seq analysis was performed by the Biomedical Informatics Shared Resource at The Ohio State University based on established protocols^23–26^. Changes in gene expression (Log_2_FC) of p<0.05 were plotted for the indicated comparisons. Details are available in online supplement. Sequence data is publicly available in the National Center for Biotechnology Information (NCBI) Gene Expression Omnibus (GEO) repository, accession no. GSE268251. Analyzed data can be found in the supplement.

### Statistical Analysis

Data is presented as mean±SEM. Sample sizes were selected based on published data to provide 80% power to detect effect sizes >2.0 at type I error α = 0.005 adjusting for multiple group comparisons. ^18,19,27^. Comparisons between two mean values were performed using the Students t-test and among more than two mean values using one-way or 2-way analysis of variance (ANOVA) or mixed-effects ANOVA (serial/repeated measures) with a Sidak or Tukey’s multiple comparison post-hoc test as indicated using Graphpad Prism 10. Statistical methods are indicated in figure legends with n and p values. Statistical significance was determined at p<0.05.

## Results

### CPT1a protein expression is selectively increased in non-ischemic cardiomyopathy

CPT1a expression increases in animal models of HF ^6, 12,27^, but recent reporting suggests that CPT1a protein is unchanged in myocardium from patients with dilated cardiomyopathy (DCM), measured via semi-quantitative proteomics ^28^. Given the potential distinctions across pathologies, we examined samples from patients with NICM from two different cohorts and different sampling protocols (collected at the University of Pennsylvania and the University of Utah), that may have pathological hypertrophy more similar to that of animal models of TAC. Myocardial CPT1 was compared between unused donor hearts and patients with NICM (Figure 1). CPT1a protein was increased in both male (Figure 1A&B) and female (Figure 1C&D) NICM patients at both institutions, with no detectable change in CPT1b. Cpt1a mRNA expression trended towards increased expression in NICM, but reached significance only in male samples from the University of Pennsylvania.

**Figure 1.**
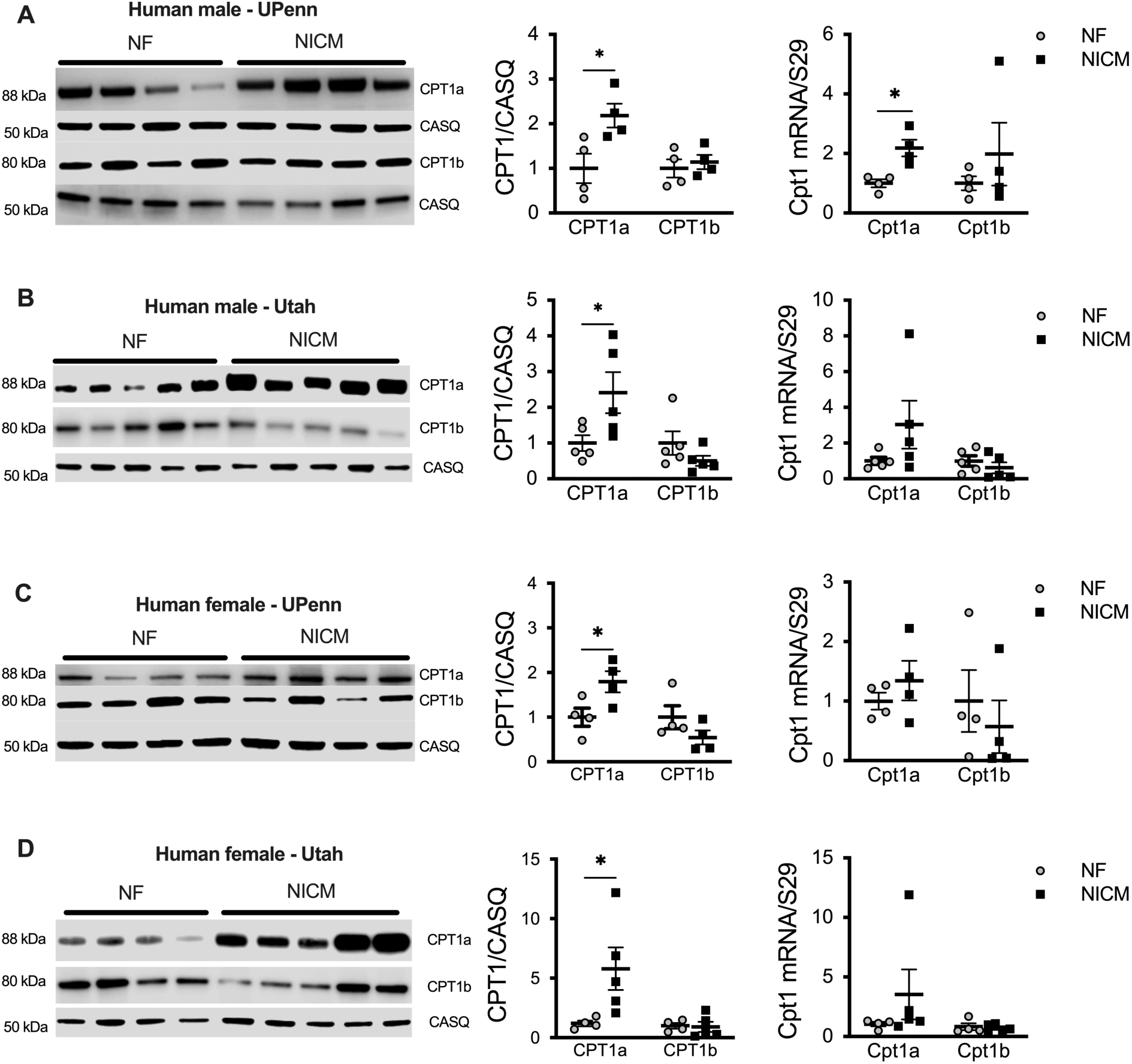
CPT1 protein and mRNA expression measured in human failing hearts from both males and females. CPT1a and CPT1b protein and mRNA (Cpt1a, Cpt1b) expression measured in male (**A&B**) and female (**C&D**) patients with non-ischemic cardiomyopathy (NICM) vs. nonfailing donor hearts (NF) from two different patient cohorts (University of Pennsylvania (UPenn), n=4 NF male and female, n=4 NICM male and female; and University of Utah (Utah), for males n=5 NF and NICM, for females n=4 NF and n=5 NICM). *p<0.05, two-way ANOVA with Sidak’s multiple comparison test.

### Increased CPT1a is an adaptive and essential response to pressure overload

To elucidate whether the CPT1a upregulation is adaptive or maladaptive in the pathologically stressed heart, cardiac specific CPT1a knockout mice (csCPT1a ko) were subjected to TAC (Figure 2A). CPT1a protein was present, but reduced 1 wk after birth in hearts of csCPT1a ko mice and absent in adult csCPT1 ko mouse hearts (Figure S2A). Mortality was increased among csCPT1a ko mice when TAC was performed with a 0.016 constriction (Figure S2B); therefore, we utilized a more moderate constriction (0.018 inches). Sham hearts of csCPT1a ko mice had significantly reduced CPT1a protein vs. f/f mice (Figure 2B), with no change in CPT1b (Figure 2B). At 8 wks post-TAC, f/f mice did not display increased CPT1a or decreased ejection fraction (EF) in response to this moderate constriction (Figure 2C). TAC did cause a 40% decrease in EF in csCPT1a ko mice (Figure 2C), despite a similar increase in LV mass from echocardiography, as in f/f littermates (Figure 2D). Heart weight to tibia length (HW:TL) measured at the end of isolated heart perfusion (discussed below) was higher in csCPT1a ko TAC vs. f/f TAC.

**Figure 2.**
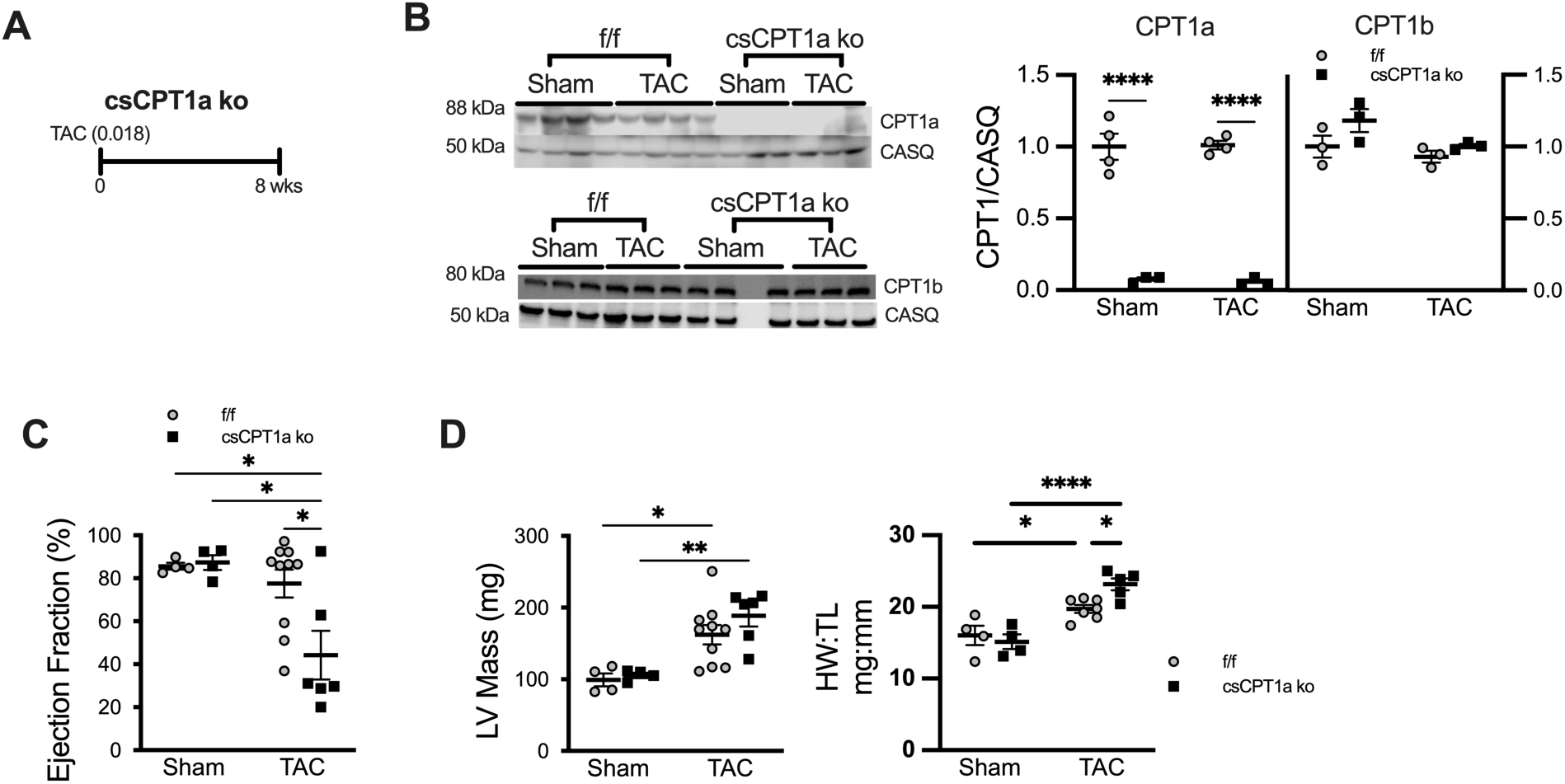
The loss of CPT1a expression sensitizes the heart to pathological stress. (**A)** Experimental protocol outlining the surgical interventions in csCPT1a ko mice. **(B)** Western blots and mean data showing CPT1a and CPT1b protein content in csCPT1a ko hearts vs. f/f 8 wks after TAC or sham surgery vs. calsequestrin (CASQ) (n=4 f/f sham, n=3 csCPT1a ko sham, n=4 f/f TAC, n=3 csCPT1a ko TAC). **(C)** Ejection fraction and left ventricular (LV) mass via echocardiography, 8 wks after TAC or sham surgery (n=4 f/f sham, n=4 csCPT1a ko sham, n=10 f/f TAC, n=6 csCPT1a ko TAC). (D) Heart weight to tibia length (HW:TL) measured in sham and TAC-operated f/f and csCPT1a ko mice at the end of isolated heart perfusion (n=4 f/f sham, n=4 csCPT1a ko sham, n=6 f/f TAC, n=7 csCPT1a ko TAC).*p<0.05, **p<0.01, ***p<0.001, ****p<0.0001, by 2-way ANOVA with Tukey’s multiple comparisons test.

How cardiac CPT1a overexpression could mediate the response to TAC was explored. AAV9.cTnT.Cpt1a delivery increased cardiac CPT1a protein (54% increase) (Figure S3A&B), but not liver (Figure S3&D) 10 d after administration (2×10^13^ viral genome/kg body weight i.v. to maintain cardiac selectivity of AAV9^29–32)^. TAC was induced in PBS or AAV9.cTnT.Cpt1a mice 10 d after i.v. injection (see Figure 3A). The increase in CPT1a was sustained in AAV9.cTnT.Cpt1a sham hearts (2.8-fold vs. PBS sham) for the 8 wk protocol duration with no compensatory changes in CPT1b (Figure 3B). Because AAV9.cTnT.Cpt1a mice did not show the increased sensitivity to TAC, as did csCPT1a ko, a more aggressive constriction was used (0.016 inch clip). CPT1a protein increased (Figure 3B), and EF and fractional shortening (FS) (Figure 3C) was reduced in hearts of PBS TAC mice with the 0.016 inch constriction. CPT1a was further augmented in hearts of AAV9.cTnT.Cpt1a TAC mice (Figure 3B) compared to PBS TAC (1.4-fold). Importantly, the declines in EF and FS (Figure 3C) at 8 wks TAC were attenuated by AAV9.cTnT.Cpt1a delivery, despite similar echocardiographic measurement of LV mass (Figure 3D). HW:TL was only increased in PBS TAC mice vs. sham (Figure 3D).

**Figure 3.**
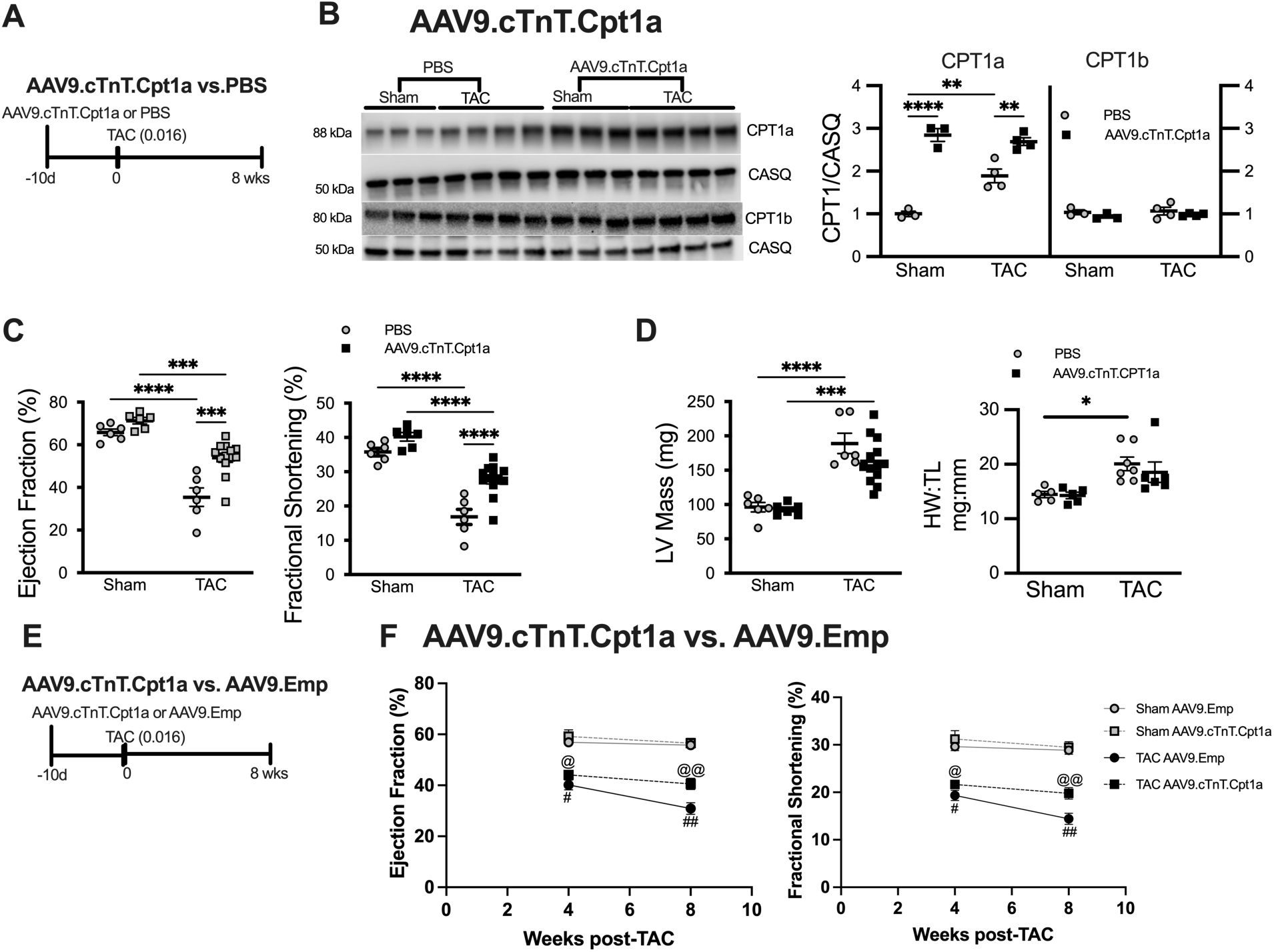
Overexpressing CPT1a in the adult mouse heart attenuates the functional decline in reponse to pathological stress. **(A)** Experimental protocol outlining the surgical interventions in C57Bl/6 mice injected with AAV9.cTnT.Cpt1a or PBS. **(B)** Western blots and mean data showing CPT1a and CPT1b protein content in hearts 8 wks after TAC or sham surgery. (**C**) CPT1a overexpression (AAV9.cTnT.Cpt1a) attenuated the reduction in ejection fraction and fractional shortening compared to PBS controls (n=6 PBS sham, AAV9.cTnT.Cpt1a sham, PBS TAC; n=13 AAV9.cTnT.Cpt1a TAC). **(D)** Left ventricular (LV) mass measured via echocardiography at 8 wks TAC (n=6 PBS sham, AAV9.cTnT.Cpt1a sham, PBS TAC; n=13 AAV9.cTnT.Cpt1a TAC) and heart weight to tibia length (HW:TL) measured at the end of isolated heart perfusion (n=6 PBS sham, n=4 AAV9.cTnT.Cpt1a sham, n=7 PBS TAC, n=6 AAV9.cTnT.Cpt1a TAC). (**E)** AAV9.cTnT.Cpt1a delivery 10 d before TAC or sham surgery similarly attenuated the decline in systolic function (**F**) when compared to empty virus treated (AAV9.Emp) (n=7 AAV9.Emp sham, n=5 AAV9.cTnT.Cpt1a sham, n=7 AAV9.Emp TAC, n=8 AAV9.cTnT.Cpt1a TAC). *p<0.05, **p<0.01, ***p<0.001, ****p<0.0001, via 2-way ANOVA with Tukey’s multiple comparisons test. @p<0.05 vs. AAV9.cTnT.Cpt1a sham 4 wks, #p<0.05 vs AAV9.Emp sham 4 wks, @@p<0.05 vs. AAV9.cTnT.Cpt1a sham 8 wks, ##p<0.05 vs. AAV9.Emp sham and AAV9.cTnT.Cpt1a TAC 8 wks, by mixed-effects ANOVA with Tukey’s multiple comparisons test.

To ensure that attenuated pathological remodeling with AAV9.cTnT.Cpt1a was not due to viral load, TAC or sham surgery was also performed in mice 10d after AAV9.cTnT.Cpt1a injection or injection of empty AAV9 (AAV9.Emp) (Figure 3E). Both the declines in EF and FS were significantly attenuated in mice receiving AAV9.cTnT.Cpt1a vs. AAV9.Emp (Figure 3F).

### FAO and CPT1a expression occur independently

In sham hearts, CPT1a deletion had no effect on the contribution of LCFA to oxidative metabolism (Figure 4). Consistent with unaltered CPT1a expression in f/f mice following less severe TAC, LCFA entry into the TCA cycle remained unchanged (Figure 4A). However, isolated heart function (Figure S4) and FAO (Figure 4A) were significantly reduced in csCPT1a ko TAC hearts.

**Figure 4.**
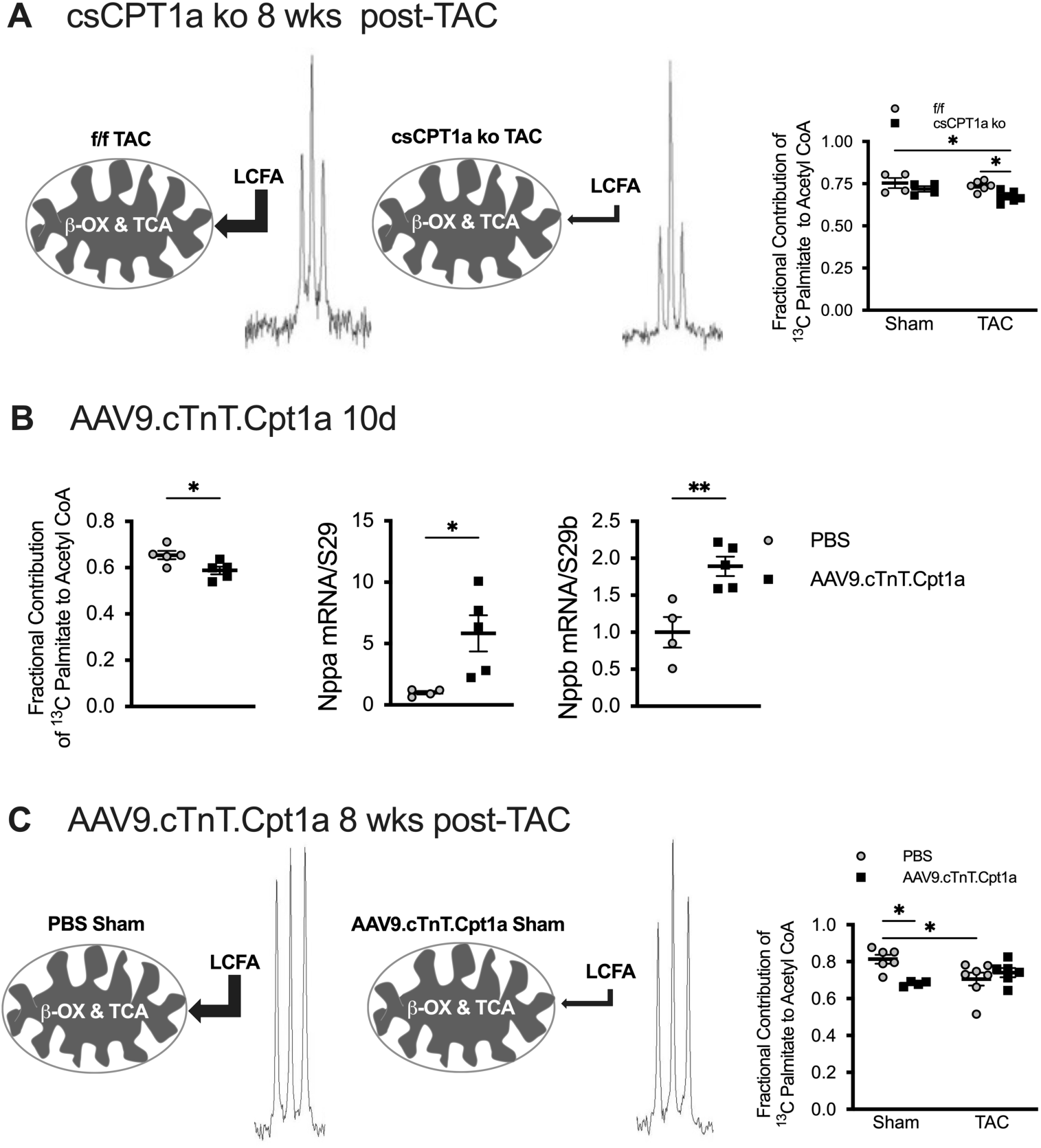
Relative long chain fatty acid oxidation measured in CPT1a knockout mice (csCPT1a ko) and mice overexpressing CPT1a (AAV9.cTnT.Cpt1a). (**A)** Representative *in vitro* ^13^C NMR signals from the 4-carbon of glutamate of f/f TAC vs. csCPT1a KO TAC hearts showing the corresponding differences in multiplet structure (singlet, glutamate labeled at the 4 carbon; doublet, glutamate labeled at the 4 carbon and the 2 or 3 carbon) of the resonance signals used to calculate the contribution of acetyl CoA derived from *β*-oxidation (*β*-OX) of long chain fatty acid (LCFA) to the tricarboxylic acid cycle (TCA) and fractional contribution of ^13^C-labeled palmitate to acetyl CoA production hearts at 8 wks after TAC or Sham surgery (n=4 f/f sham, n=4 csCPT1a ko sham, n=6 f/f TAC, n=7 csCPT1a ko TAC). **(B)** Fractional contribution of ^13^C palmitate to acetyl CoA production 10 d after AAV9.cTnT.Cpt1a delivery (n=5) vs. PBS (n=5) and cardiac ANP (Nppa) and BNP (Nppb) mRNA expression were increased 10 d after AAV9.cTnT.Cpt1a delivery (n=5) vs PBS (n=5). **(C)** Representative NMR signals from the 4-carbon of glutamate from AAV9.cTnT.Cpt1a vs. PBS control hearts used to calculate the contribution of acetyl CoA derived from *β*-oxidation (*β*-OX) of long chain fatty acid (LCFA) to the tricarboxylic acid cycle (TCA) and contribution of ^13^C palmitate to acetyl CoA production (n=6 PBS sham, n=4 AAV9.cTnT.Cpt1a sham, n=7 PBS TAC, n=6 AAV9.cTnT.Cpt1a TAC). For A and C, *p<0.05, by 2-way ANOVA with Tukey’s multiple comparisons test. For B, *p<0.05, unpaired 2-tailed t-test.

Acutely increasing CPT1a led to reduced LCFA entry into the TCA cycle (Figure 4B) and an increase in ANP (Nppa) and BNP (Nppb) gene expression prior to any surgical intervention, consistent with previous observations of acute delivery of CPT1a via adenovirus ^12^. Entry of LCFA into the TCA cycle (Figure 4C) was also lower in AAV9.cTnT.Cpt1a sham hearts, but there was no TAC-induced decline in LCFA contribution to the TCA cycle, as otherwise occurred in PBS TAC hearts (Figure 4C). Importantly, there was no difference in the contribution of LCFA to oxidation in the TCA cycle between PBS TAC and AAV9.cTnT.Cpt1a TAC, despite the attenuation of pathological remodeling in AAV9.cTnT.Cpt1a TAC.

Isolated heart function was significantly reduced in both PBS TAC and AAV9.cTnT.Cpt1a TAC hearts (Figure S5) vs. sham. Although mean isolated heart function trended strongly toward increased function in AAV9.cTnT.Cpt1a TAC vs. PBS TAC, observed increases in mean rate pressure product (RPP) and rates of pressure development (+dp/dt) and relaxation (-dp/dt) did not reach statistical significance via 2-way ANOVA. Mean RPP and dp/dt were significantly higher for AAV9.cTnT.Cpt1a TAC hearts in one-to-one comparison to PBS TAC hearts (Students t-test).

### Upstream and endogenous microRNA regulation of cardiac CPT1a expression in response to pathological stress

MiR370 suppresses hepatic Cpt1a expression ^17^, but regulation of cardiac CPT1a by miR370 and the response to cardiac stress has not been previously investigated. MiR370 content was significantly reduced in failing hearts of both mice (Figure 5A) and NICM patients (Figure 5B), suggesting a regulatory mechanism for increasing CPT1a in both animal models of HF and human NICM (Figure 1). Overexpressing miR370 in the heart (i.v. AAV9.cTNT.miR370) suppressed Cpt1a expression without affecting Cpt1b (Figure 5C). AAV9.cTnT.miR370 did not alter the expression of miR370 in the liver or skeletal muscle (Figure S6). Delivering AAV9.cTnT.miR370 to mice 10d prior to TAC (Figure 5D) suppressed upregulation of CPT1a by TAC, but CPT1a expression remained at 50% of PBS control (Figure 5D). Cardiac function was equally suppressed in mice with TAC following PBS or miR370 delivery (Figure 5E), and the TAC-induced decrease in FAO was unaffected (Figure 5F). Thus, the decline in FAO in response to TAC can occur independently of the upregulation of CPT1a, but a requisite level of CPT1a attenuates the increased sensitivity of csCPT1a ko mice to TAC. Thus, increased CPT1a in pathologically stressed hearts is facilitated through reduced miR370 expression.

**Figure 5.**
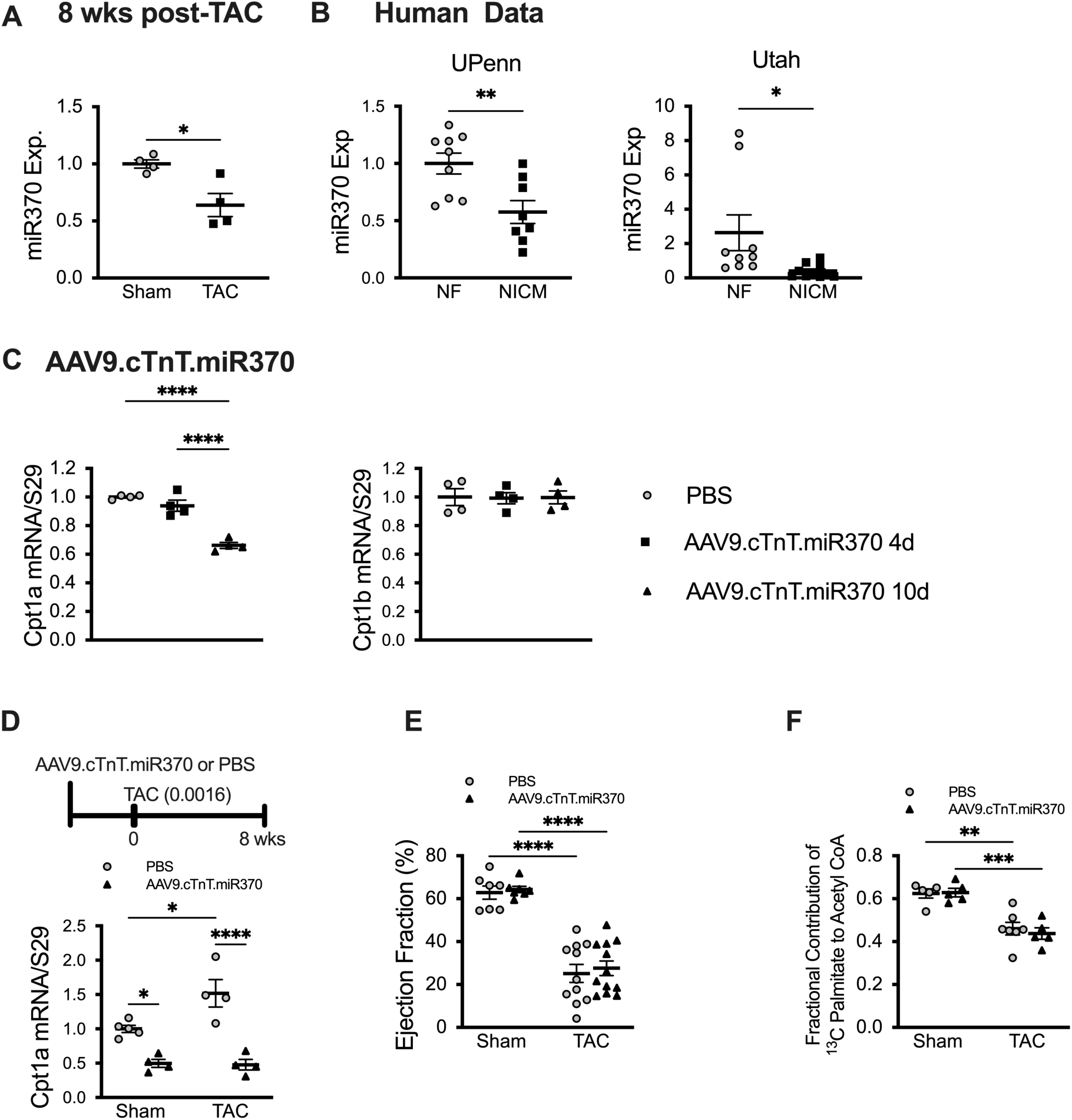
MiR370 regulates CPT1a expression in heart failure. **(A)** MiR370 expression in mice 8 wks after transverse aortic constriction (TAC) or sham surgery (n=4). **(B)** MiR370 expression in non-failing (NF) heart tissue and heart tissue from patients with non-ischemic cardiomyopathy (NICM) collected at the University of Pennsylvania (n=4 male and n=4 female combined for both NF and NICM) and the University of Utah (n=5 male and n=4 female NF, n=5 male and female NICM). **(C)** Cpt1a and Cpt1b mRNA expression in the heart following i.v. AAV9.cTnT.miR370 measured at the indicated time points (n=4; one-way ANOVA with Tukey’s multiple comparisons test). **(D)** Depiction of experimental protocol for mir370 delivery followed by TAC and Cpt1a mRNA expression 8 wks after TAC or sham surgery in mice previously injected with PBS or AAV9.cTnT.miR370 (n=5 PBS sham, n=4 AAV9.cTnT.miR370 sham, n=4 PBS TAC, n=4 AAV9.cTnT.miR370 TAC). Ejection fraction **(E)** (n=7 PBS sham, n=7 AAV9.cTnT.miR370 sham, n=11 PBS TAC, n=12 miR370 TAC) and fractional contribution (n=5 PBS sham, n=5 AAV9.cTnT.miR370 sham, n=7 PBS TAC, n=5 AAV9.cTnT.miR370 TAC) of ^13^C palmitate to acetyl CoA production **(F)** in PBS or AAV9.cTnT.miR370 mice 8 wks after TAC or sham surgery. *p<0.05, **p<0.01, ***p<0.001, ****p<0.0001; unpaired 2-tailed t-test (A&B), one-way ANOVA with Tukey’s multiple comparisons test (C), and 2-way ANOVA with Tukey’s multiple comparisons test (D-F).

### Elevated ANP expression in response to CPT1a overexpression is transient

Acutely increasing CPT1a (via AAV9.cTnT.Cpt1a) led to increased cardiac Nppa expression (Figure 4C), as observed previously with adenovirus ^12^. Increased Nppa expression was a selective effect of CPT1a overexpression, as AAV9.Emp (Figure S7) and AAV9.cTnT.miR370 did not alter Nppa (Figure 6A). Strikingly, by 8 wks post-surgery, upregulation of Nppa was no longer evident in AAV9.cTnT.Cpt1a TAC or sham mice (Figure 6B). Conversely, TAC increased Nppa expression in PBS treated and csCPT1a ko mice (Figure 6B).

**Figure 6.**
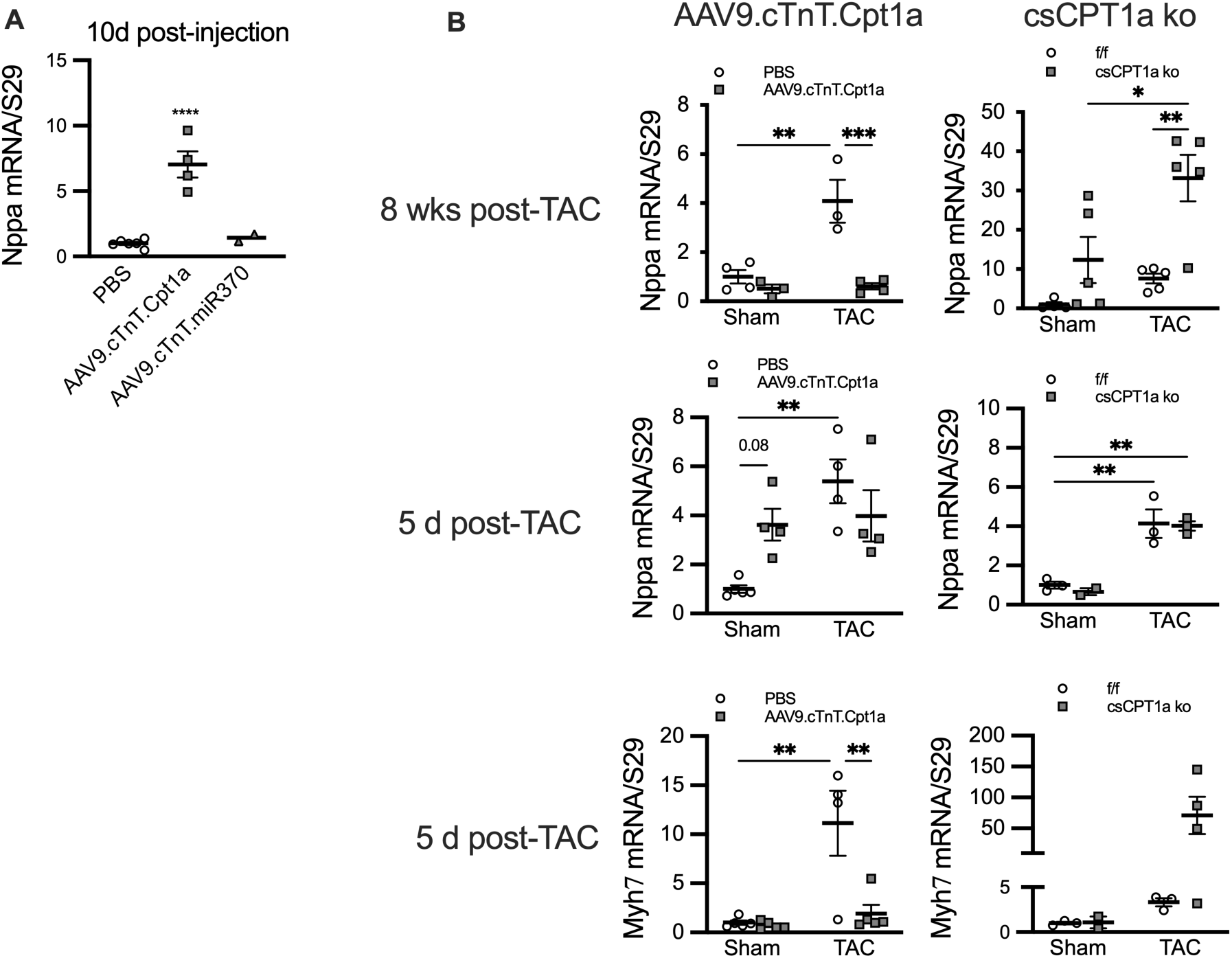
ANP mRNA (Nppa) Expression in Response to TAC in cardiac specific CPT1a knockout mice (csCPT1a ko) and mice overexpressing CPT1a (AAV9.cTnT.Cpt1a). **(A)** Cardiac Nppa expression 10 d after injection of AAV9.cTnT.Cpt1a or AAV9.cTnT.miR370 (n=6 PBS, n=4 AAV9.cTnT.Cpt1a, n=2 AAV9.cTnT.miR370). (**B)** Cardiac Nppa and Myh7 expression measured at the indicated time points <u>in</u> AAV9.cTnT.Cpt1a (0.016 clip) and csCPT1a ko mice (0.018 clip) (8 wks AAV9.cTnT.Cpt1a, n=4 PBS sham, n=3 PBS TAC, n=3 AAV9.cTnT.Cpt1a sham, n=4 AAV9.cTnT.Cpt1a TAC; 8 wks csCPT1a ko, n=5; 5 d AAV9.cTnT.Cpt1a, n=5 PBS sham, all others n=4; 5 d csCPT1a ko, n=3). *p<0.05, **p<0.01, ***p<0.001, <u>one-way ANOVA</u> (A) and 2-way ANOVA with Tukey’s multiple comparisons test (B).

To distinguish the transient increase in Nppa expression in response to CPT1a overexpression from the TAC-induced response in Nppa, we examined hearts 5 d after TAC (Figure 6B). Results were compared to csCPT1a ko mice at the same time point using the TAC protocols depicted in Figure 2A for csCPT1a ko mice and AAV9.cTnT.Cpt1a mice. AAV9.cTnT.Cpt1a sham hearts showed a strong trend toward increased Nppa expression vs. PBS sham at 5 d (Figure 6B). However, TAC did not lead to a further increase in Nppa; while in f/f and csCPT1a ko hearts, there was no difference in Nppa expression during the acute response to TAC. In AAV9.cTnT.Cpt1a treated mice, the priming of Nppa upregulation in advance of TAC was associated with attenuated hypertrophic remodeling at this early time point, delaying upregulation of *β*-myosin heavy chain mRNA (Myh7) (Figure 6B), while in csCPT1a ko mice there was a strong trend towards increased Myh7 expression (Figure 6B) at this early time point. By 4 wks TAC Nppa expression was no longer increased in AAV9.cTnT.Cpt1a treated mice (Figure S7A), and Myh7 expression (Figure S7B) was not different between PBS and AAV9.cTnT.Cpt1a treated mice.

### Intervening with CPT1a overexpression after onset of cardiac dysfunction during pressure overload mitigates adverse remodeling and functional decline

To determine if CPT1a-induced upregulation of Nppa is required prior to pathological stress to attenuate adverse remodeling, the therapeutic potential of increasing CPT1a post-TAC after the onset of dysfunction was investigated (Figure 7A). There was a significant reduction in EF and FS (Figure 7B) prior to AAV9 gene delivery at 4 wks after TAC or sham surgery, which were rescued by AAV9.cTnT.Cpt1a administration at both 2 and 4 wks after injection without affecting LV mass (Figure 7C). CPT1a protein was increased in both sham and TAC mice receiving AAV9.cTnT.Cpt1a vs. AAV9.Emp (Figure 7D). Predictably, Nppa expression was increased in AAV9.Emp TAC (Figure 7E). In contrast, AAV9.cTnT.Cpt1a TAC mice did not have increased Nppa. Both AAV9.cTnT.Cpt1a and AAV9.Emp TAC hearts did have elevated Myh7 expression at 8 wks post-TAC (Figure 7E).

**Figure 7.**
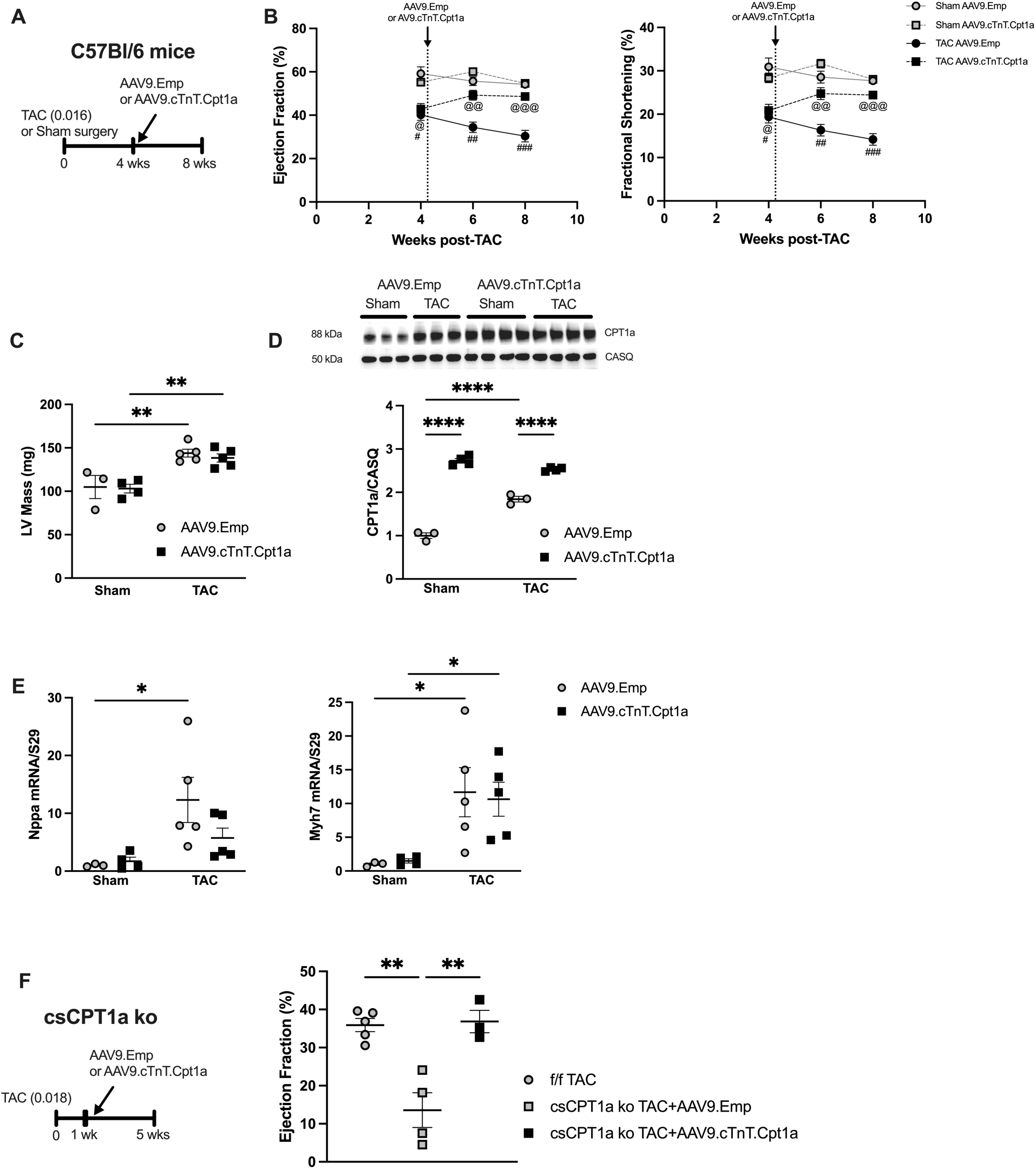
Increasing CPT1a expression exhibits therapeutic potential. **(A)** AAV9.cTnT.Cpt1a or AAV9.Emp was administered 4 wks after TAC or sham surgery in C57Bl/6 mice. **(B)** The decline in ejection fraction and fractional shortening was partially corrected 2 and 4 wks after AAV9.cTnT.Cpt1a administration (n=3 Sham AAV9.Emp, n=4 sham AAV9.cTnT.Cpt1a, n=5 TAC AAV9.Emp, n=5 TAC AAV9.cTnT.Cpt1a). **(C)** LV mass measured via echocardiography 8 wks post-TAC in C57Bl/6 mice described in **(A)**. CPT1a expression **(D)** as well as **(E)** ANP (Nppa) and *β* myosin heavy chain mRNA (Myh7) measured at the end of the protocol described in **(A)**. **(F)** Treatment of csCPT1a ko mice post-TAC with AAV9.cTnT.Cpt1a or AAV9.Emp, 1 wk after TAC surgery and ejection fraction measured at the end of the treatment protocol described in (n=5 f/f TAC, n=4 csCPT1a ko+AAV9.cTnT.Cpt1a TAC, n=3 csCPT1a ko+AAV9.Emp TAC). #p<0.05 vs Sham AAV9.Emp at 4 wks, ##p<0.05 vs. Sham AAV9.Emp and TAC AAV9.cTnT.Cpt1a at 6 wks, ###p<0.05 vs. Sham AAV9.Emp and TAC AAV9.cTnT.Cpt1a at 6 wks, @ p<0.05 vs Sham AAV9.cTnT.Cpt1a at 4 wks, @@p<0.05 vs Sham AAV9.cTnT.Cpt1a at 6 wks, @@@p<0.05 vs Sham AAV9.cTnT.Cpt1a at 8 wks; by mixed-effects ANOVA with Tukey’s multiple comparisons test (B). *p<0.05, **p<0.01, ***p<0.001, ****p<0.0001; by 2-way ANOVA with Tukey’s multiple comparisons test (C-F).

In addition to demonstrating therapeutic potential in wild type mice, rescue with AAV9.cTnT.Cpt1a treatment was also effective in csCPT1a ko mice after TAC. CsCPT1a ko mice underwent TAC surgery and were given AAV9.cTnT.Cpt1a or AAV9.Emp 1 week later (Figure 7F). Introducing CPT1a into the hearts of csCPT1a ko mice significantly attenuated the dramatic decline in EF otherwise occurring in csCPT1a ko mice treated with empty virus (Figure 7F). Therefore, CPT1a overexpression prior to pathological stress is not required to exert cardioprotective effects, and can effectively be used therapeutically to mitigate adverse remodeling after the onset of dysfunction.

### The expression of CPT1a regulates cardiac gene expression

In sham hearts, csCPT1a knockdown led to significant changes in expression of 1328 genes, primarily inducing expression, while CPT1a overexpression led to changes in expression of 852 genes, primarily suppressing expression (Figure S10). A number of genes suppressed by AAV9.cTnT.Cpt1a in sham hearts were highly enriched in pathways involved in fibrosis, cell growth and apoptosis (Table S6).

Figure 8A shows the overlap in genes affected by both CPT1a knockdown and overexpression in sham hearts, and the change in expression (log_2_FC) for each of those commonly affected genes is plotted in Figure 8B. Pathway enrichment analysis of these common genes indicated enriched apoptotic pathways, profibrotic-related pathways, and the response to TGFb; they also extend to pathways involved in cell cycle regulation and lipid metabolism (Figure 8C). Therefore, in the absence of pathological stress, changes in CPT1a expression significantly alters gene expression patterns. A recent study of CPT1b knockdown suggested upregulation of proliferative pathways in the heart ^33^. Although histochemistry revealed a trend towards increased cardiomyocytes with PHH3 positive nuclei 10 d after AAV9.cTnT.Cpt1a injection, this did not reach full statistical significance (p=0.054, Figure S7).

**Figure 8.**
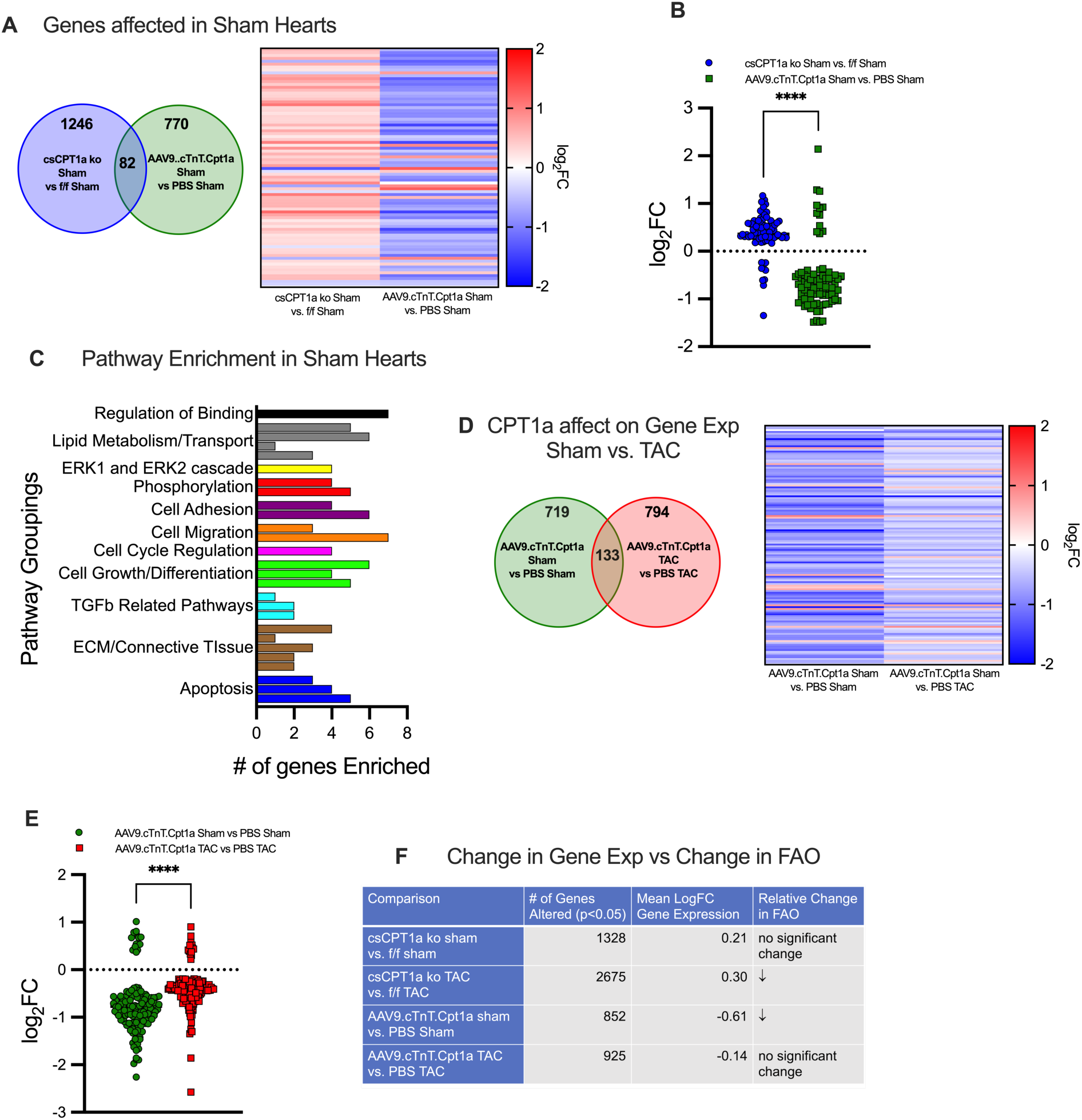
CPT1a expression changes lead to profound differences in overall gene expression. **(A)** Diagram showing the number of genes altered by cardiac specific CPT1a knockout (csCPT1a ko, n=4) or cardiac specific CPT1a overexpression (AAV9.cTnT.Cpt1a, n=3 PBS sham, n=4 AAV9.cTnT.Cpt1a sham) and heatmap of changes in gene expression that occurred in the 82 overlapping genes that were significantly altered by both the loss of CPT1a and AAV9.cTnT.Cpt1a. **(B)** Mean change in gene expression (log_2_FC) for the 82 overlapping genes altered in **(A)**. **(C)** Pathway enrichment for the 82 overlapping genes identified in **(B)**. The number of genes enriched in each pathway is indicated and related pathways were grouped together. **(D)** Diagram of genes altered by AAV9.cTnT.Cpt1a in either sham or TAC hearts (8 wks post-TAC) and heatmap for the 133 overlapping genes for which the expression was changed by CPT1a overexpression vs. both PBS sham and PBS TAC. **(E)** Mean change in gene expression (log_2_FC) for those 133 overlapping genes altered in **(C)**. **(F)** Summary of the changes in gene expression and corresponding changes in LCFA contribution to oxidative metabolism (FAO) revealing independent responses to altered CPT1a expression for the comparisons indicated. ****p<0.0001; by unpaired 2-tailed t-test.

In TAC hearts, the inverse relationship between CPT1a expression and gene expression patterns was maintained. Figure 8D shows overlap between patterns induced by CPT1a overexpression in both sham (AAV9.cTnT.Cpt1a sham vs. PBS sham) and TAC hearts (AAV9.cTnT.Cpt1a TAC vs. PBS TAC). Overall, 133 genes were altered by AAV9.cTnT.Cpt1a under both conditions (Figure 8D,E). AAV9.cTnT.Cpt1a delivery increased CPT1a 82% in sham hearts (AAV9.cTnT.Cpt1a vs. PBS sham) but only a 42% in TAC hearts ((AAV9.cTnT.Cpt1a TAC vs. PBS TAC), as CPT1a was induced by TAC itself (Figure 2). Therefore, the relative reduction in gene expression induced by AAV9.cTnT.Cpt1a (Figure 8E) was proportional to the relative change in CPT1a. FAO was either unchanged or reduced when comparing the indicated groups in Figure 8F and did not correlate to the change in CPT1a expression. Importantly, combined results from Sham and TAC hearts showing similar effects of CPT1a expression on gene suppression demonstrate that attenuated gene programs for adverse changes in TAC hearts overexpressing CPT1a are not merely due to improved cardiac health. Indeed, when considered with the induction of these adverse gene programs in csCPT1a KO hearts, the overall results substantiate the beneficial effects of CPT1a expression on suppression of adverse gene programs in the failing heart.

Based on RNAseq results, expression of collagen markers was assessed in TAC hearts from both AAV9 treated and csCPT1a ko mice (Figure S11). Collagen type III alpha 1 (Col3a1) was selectively upregulated in models with reduced CPT1a expression 8 wks after TAC, AAV9.cTnI.miR370 TAC hearts and csCPT1a ko TAC hearts (Figure S11A&B). In hearts from C57Bl/6 mice treated with AAV9.cTnT.Cpt1at 4 wks TAC there was a significant reduction in both Col3a1 and Tgfb expression vs. TAC AAV9.Emp treated mice (Figure S11C).

## Discussion

This is the first study to evaluate the role for, and impact of CPT1a upregulation in the development of HF. The data are consistent with previously reported increases in cardiac CPT1a in animal models of HF ^27^, and clearly demonstrate CPT1a upregulation in male and female patients with NICM. This study elucidates upregulation of CPT1a as a critical, cardioprotective response to pathological stress. The absence of CPT1a sensitizes the heart to afterload stress, leading to accelerated decompensation and LV dilation, while overexpressing CPT1a attenuates the progression of decompensatory hypertrophy. The innate stress-induced increase in CPT1a, partially mediated through a reduction in miR370 expression, is insufficient, and augmenting CPT1a content holds significant therapeutic potential. Importantly, cardioprotection was realized with AAV9.cTnT.Cpt1a administration even at the point of decompensated hypertrophy and dysfunction in C57/Bl6 and csCPT1a ko mice. CPT1a expression induces suppression of gene expression programs, including but not limited to those involved in pathological remodeling. Similar gene suppression via CPT1a was observed in both sham and TAC hearts receiving AAV9.cTNT.CPT1a, while csCPT1a KO hearts displayed induction of these same genes irrespective of sham or TAC status. Therefore, the altered gene expression profiles of hearts receiving AAV9.cTNT.CPT1a during TAC are not the mere consequence of other beneficial effects of CPT1a expression, but result from the direct effects of CPT1a on gene expression.

Increasing *in vivo* CPT1a expression in adult hearts, whether via adenoviral vector ^12^ or AAV9.cTnT.Cpt1a administration (Figure 3), leads to reduced cardiac FAO, without any change in mechanical work. In adult hearts, CTP1b is the predominant isoform, while developing hearts express CPT1a at relatively high levels ^3,4^. Baseline CPT1a in the adult heart is low, estimated to contribute less than 10% to total activity ^3^. *Ex vivo* tissue studies report that negative feedback inhibition of CPT1b occurs by allosteric regulation by malonyl CoA ^13^, but CPT1a was shown to be less sensitive to malonyl CoA inhibition in non-cardiac cells ^13^, Thus, it may be surprising that increased CPT1a in cardiomyocytes causes reduced FAO, yet this is previously reported and consistent with disassociation between cardiac malonyl CoA content and FAO ^12, 15,16,34^.

Upregulation of CPT1a in HF is a reversal of the shift in cardiac CPT1 isoforms that occurs during development from neonate to adult. Although sometimes argued that CPT1b is lower in fetal hearts^35^, CPT1b content is consistent throughout development, while CPT1a decreases ^3,36^. We reconfirmed that CPT1b protein is not altered in failing hearts of mice and humans. Thus, if CPT1a is associated with reduced FAO, which persisted in the present study throughout prolonged CPT1a overexpression in adult hearts, then the decline in CPT1a in the normal adult heart may serve to activate overall CPT1 activity. Surprisingly, decreasing CPT1a in the present study did not alter baseline FAO. This indicates that reducing CPT1a expression beyond its inherently low level, as observed with CPT1a knockdown and miR370 overexpression, does not further augment FAO because the action of CPT1b on FAO is already heightened.

In hearts overexpressing CPT1a, TAC did not induce further reduction of FAO vs. sham, and FAO was not significantly different vs. PBS TAC. This is not surprising as there was no increase in CPT1a expression between sham and TAC hearts overexpressing CPT1a, and a smaller difference in CPT1a expression between TAC hearts as was observed between shams. Indeed, in tissues with high CPT1a, CPT1a content is not proportional to FAO rate ^14^.

Myocardial CPT1a protein was increased in NICM patients. CPT1a mRNA expression has been reported to be unchanged in human HF ^35,37,38^, but protein expression has not been widely reported. However, as shown here, mRNA and protein cannot be used interchangeably. A recent report did not note increased CPT1a in patients with DCM using a semi-quantitative proteomics-based method ^28^. The differing results could reflect the nature of analysis or the heterogeneity of HF, even among patients with HFrEF. Significantly, CPT1a was increased in myocardium from separate cohorts of NICM patients at two different institutions.

Substrate labelling studies in HF patients are consistent with reduced FAO ^10,11^, and could be a consequence of not only altered mitochondrial fatty acyl CoA entry, but also a reduction in fatty acid uptake and activation in the heart ^18^. Irrespective of these differences in past observations, the most salient findings of the present study are not the degree to which CPT1a is increased in failing hearts, but rather the adaptive and essential nature of that increase, and the therapeutic potential of increasing CPT1a expression in the failing heart.

Increased myocardial CPT1a in TAC mice and NICM patients is associated with reduced miR370 expression. In liver, miR370 regulates CPT1a expression^17^, and this is the first report of miR370 regulating cardiac CPT1a expression in normal and pathological hearts. Overexpressing miR370 prevented the TAC-induced increase in CPT1a, but did not greatly sensitize the heart to pathological stress. MiR370 has been shown to have effects beyond CPT1a regulation ^39^, but it is important to consider that CPT1a expression was reduced, not absent in AAV9.cTnT.miR370 hearts.

Acutely increasing CPT1a induces increased ANP expression, and although ANP is often viewed as a marker of the cardiac response to pressure overload and pathological hypertrophy, previous studies showed that ANP attenuates pathological remodeling and blocking ANP expression accelerates cardiac decompensation ^40,41^. Perhaps ANP upregulation by CPT1a is participatory in attenuating cardiac remodeling in response to TAC. However, this increased ANP expression was not maintained beyond 8 weeks, and ANP induction prior to pathological stress was not requisite for the beneficial effects of CPT1a.

The significant impact that CPT1a expression has on gene regulatory pathways and the adaptive response to chronic pressure overload is striking. CPT1a has been identified as a key regulator of gene expression in cancer cells, with increased CPT1a expression resulting in increased acetylation and tumor progression through inhibition of apoptosis ^42^. The regulation of gene expression by CPT1a is documented to occur independently of FAO in some cancer cells ^43^. Braun, *et al*, reported suppression of CPT1b in cardiomyocytes led to upregulation of a proliferative gene expression pattern ^33^. While they reported no change in CPT1a with reduced CPT1b, the isoform ratio may have been altered. Just as for FAO, the gene expression patterns of the heart may be sensitive to the ratio of CPT1a to CPT1b.

CPT1a is shown here to actively contribute to suppression of pathways involved in fibrosis. While a reduced profibrotic gene program could be attributed to noncardiac cells secondary to improved outcomes due to CPT1a overexpression in the pathological heart, CPT1a also affected fibrotic genes in sham hearts in which there is no alleviation of pathology. Indeed, Col3a1 in particular was suppressed by AAV9.cTNT.CPT1a in TAC hearts, and Col3a1 expression is documented in cardiomyocytes ^44^, while both CPT1a knockdown and miR370 overexpression increased Col3a1. Furthermore, Tgfb has a cardiomyocyte component in addition to fibroblasts ^45,46^, and cardiomyocytes can influence collagen formation indirectly ^47^. Thus, the effects of CPT1a on fibrotic gene program expression are unlikely to be due to non-cardiac cell responses alone, and the evidence suggests a direct effect of CPT1a expression within the targeted cardiomyocytes.

Based on these novel findings, increased CPT1a expression and content play an essential role in the cardiac stress response, mediating cardiac metabolic and functional remodeling and regulating stress-induced cardiac gene expression. Importantly, the translational relevance of these findings is that overexpressing CPT1a following TAC, even after emergence of cardiac dysfunction, reversed the decline in EF and attenuated HFrEF, and that even the csCPT1a ko mouse can be partially rescued after the induction of pathological stress through re-expression of CPT1a. CPT1a in the adult heart has an essential role in regulating gene programs (Figure 8), even absent pathological stress, with the capacity of the heart to both respond to pathological stress and regulate cardiac gene expression proportional to the induction of CPT1a expression. The changes in gene expression do not correlate with a specific change in FAO and reveal a previously unknown non-canonical role of CPT1a in responding to pathological stress on the heart.

## Non-standard Abbreviations and Acronyms

AAV: adeno-associated virus
AAV9: adeno-associated virus serotype 9
αMHC: α-myosin heavy chain
ANP: atrial natriuretic peptide
CASQ: calsequestrin
CF-LVAD: continuous-flow left ventricular assist device
cTNT: cardiac troponin T
Col1a1: collagen type I alpha 1 chain gene
Col3a1: collagen type III alpha 1 chain gene
CPT1: carnitine palmitoyl transferase 1 cs
CPT1a ko: cardiac specific CPT1a knockout
DCM: dilated cardiomyopathy
EF: ejection fraction
FAO: long chain fatty acid oxidation
FS: fractional shortening
GAPDH: glyceraldehyde 3-phosphate dehydrogenase
HF: heart failure
HFrEF: heart failure with reduced ejection fraction
LCFA: long chain fatty acid
LV: left ventricular
miR370: microRNA 370
Myh7: β-myosin heavy chain gene
NICM: nonischemic cardiomyopathy
NP: natriuretic peptide
Nppa: natriuretic peptide A gene
NPPb: natriuretic peptide B gene
TAC: transverse aortic constriction
TGFβ: transforming growth factor β

## Acknowledgements

The authors are grateful to the donor families for their generosity along with DonorConnect of Salt Lake City, UT and the Gift of Life Donor Program of Philadelphia, PA for facilitating the procurement of the human myocardial tissue by our research teams.

## Sources of Funding

This work was supported by funding from the National Heart Lung and Blood institute (NHLBI) of the National Institutes of Health (NIH) grants R01HL132525 (E.D.L.), R01HL160646 (E.D.L.), R01HL164290 (E.D.L.), R01HL153876 (H.Z), R01HL156667 (S.D.), R01HL135121 (S.D.), P30 CA016058 (A.W.), as well as AHA 16SFRN29020000 (S.D.), and the Nora Eccles Treadwell Foundation (S.D.). A.A.Z. was supported by NIH training grant T32HL149637 and C.P.K. was supported by NIH training grant T32HL007576.

## Disclosures

None

## Novelty and Significance

### What is known?

- CPT1a protein expression is increased in myocardium of preclinical animal models of heart failure (HF), coinciding with reduced fatty acid oxidation (FAO).
- Acutely increasing CPT1a in the heart is associated with a decrease in FAO and increased natriuretic peptide expression.
- Whether the increase in CPT1a in HF is adaptive or maladaptive is unknown.

### What new information does the article contribute?

- CPT1a protein expression is increased in myocardium of male and female patients with HF, and this shift in cardiac CPT1 isoforms is inverse to the shift in miR370 expression, consistent with miR370 regulation of CPT1a in mouse hearts.
- CPT1a upregulation in response to pathological stress is an essential adaptive response, and increasing CPT1a expression in HF significantly mitigates adverse pathological remodeling.
- Independent of effects on fatty acid oxidation, CPT1a expression induces the suppression of gene expression programs including but not limited to those involved in pathological remodeling.

CPT1a is shown to be a potential therapeutic target for HF that rescues contractile function when overexpressed after hypertrophic decompensation. The beneficial effects of cardiac specific, CPT1a gene delivery to failing hearts holds potential as a gene therapy approach to mitigate adverse pathological remodeling that leads to overt HF.

## Supplemental Material

Supplemental Methods

Tables S1–S6

Figure S1-S11

Major Resources Table

References #–#

Analyzed RNA-seq Data

## Notes

### Competing Interest Statement

The authors have declared no competing interest.

